# *Cyniclomyces guttulatus* is an opportunistic pathogen in rabbits with coccidiosis

**DOI:** 10.1101/850230

**Authors:** Tuanyuan Shi, Hongchao Sun, Yuan Fu, Hao Lili, Yongxue Zhou, Liu Yan, Guolian Bao, Xun Suo

## Abstract

*Cyniclomyces guttulatus* is a common inhabitant of the gastrointestinal tract in rabbits, and large numbers are often present in feces of diarrheic rabbits. However, its relation with rabbit diarrhea has not been clearly identified. We isolated a *C. guttulatus* Zhejiang strain from a rabbit with severe diarrhea and optimized the culture conditions in YPG medium. The sequenced 18S and 26S ribosomal DNA fragments were 1559bp and 632bp, respectively, and showed 99.8% homology with the 18S ribosomal sequence of the NRRL Y-17561 isolate from the dog and 100% homology with the 26S ribosomal sequence of the DPA-CGR1 and CGDPA-GP1 isolates from the rabbit and guinea pig. Our isolate was not pathogenic to healthy SPF rabbits. Instead, rabbits inoculated with the yeast had a slightly better body weight gain and higher food intake. Rabbits co-inoculated with *C. guttulatus* and the coccidian, *E. intestinalis* developed more severe coccidiosis as shown by clinical signs, and decreased body weight gain, diarrhea and death, associated with significantly higher fecal output of *C. guttulatus* vegetative cells but lower coccidian oocysts output than the rabbits inoculated with *C. guttulatus* or *E. intestinalis* alone. We also surveyed the prevalence of *C. guttulatus* in rabbits and found a positive rate of 83% in Zhejiang province. Our results indicate that *C. guttulatus* alone is not pathogenic to healthy rabbits, but could become an opportunistic pathogen when the digestive tract is damaged by other pathogens such as coccidia.

**Author summary:** *Cyniclomyces guttulatus*, a commensal yeast in rabbit gastrointestinal tract, is very commonly seen in diarrhea cases. However, it is unclear whether it causes or is a co-cause of diarrhea with other pathogens. Here, a *C. guttulatus* Zhejiang strain was firstly isolated from a rabbit with severe diarrhea and the culture conditions in YPG medium were optimized. Then, it was identified in morphology and molecular. It was agreed with the previous description in morphology and showed a closer phylogenetic relationship with other strains originated from herbivores than those from the carnivore. Finally, the *C. guttulatus* Zhejiang strain was inoculated to SPF rabbits singly or co-inoculated with *Eimeria intestinalis*. All of the results in animal assays show *C. guttulatus* alone is not pathogenic but seems a probiotic microorganism in rabbits. However, it could become an opportunistic pathogen when the digestive tract is damaged by other pathogens such as coccidia.

## Background

Diarrhea is very common in rabbits, especially in weanling rabbits, and causes huge losses to rabbitry. According to some reports, more than 50% of mortalities in rabbits could be caused by intestinal disease with diarrhea^[1–3]^. Currently, more than twenty microorganisms including viruses (e.g. *Lapine rotavirus*), bacteria (e.g. *Esherichia coli, Salmonella typhimurium, Clostridium welchii* and *Pasteurella multocida*) and parasites (e.g. *Eimeria spp., Passalurus ambiguus, Cryptosporidium spp*. and *Giardia duodenalis*) have been identified as pathogens causing diarrhea in rabbits^[4–15]^. In addition to the above pathogens, *Cyniclomyces guttulatus*, a commensal yeast in rabbit gastrointestinal tract is also commonly seen in diarrhea cases. However, it is unclear whether it causes or is a co-cause of diarrhea with other pathogens. Some researchers believed that *C. guttulatus* was not a pathogen causing diarrhea, but is probably a salubrious normal inhabitant based on its common existence in healthy animals and the absence of clinical signs in experimental rabbits inoculated with *C. guttulatus* isolates^[16,17]^. However, other researchers believed it could be an opportunistic pathogen based on the large number of yeast cells in feces of diarrheic animals and the positive response of some diarrheic cases to anti-fungal treatment with nystatin^[18–22]^. To clearly establish the relationship between *C. guttulatus* and diarrhea in rabbits, a *C. guttulatus* strain was isolated and identified from a rabbit with severe diarrhea. Then, its relationship to rabbit diarrhea was investigated through inoculation of *C. guttulatus* alone and co-inoculation with an intestinal protozoan, *Eimeria intestinalis*. In addition, the prevalence of *C. guttulatus* in rabbits was surveyed in Zhejiang province of China.

## Materials and methods

### Isolation and cultivation of *C. guttulatus*

A *C. guttulatus* Zhejiang strain was isolated from a severely diarrheic rabbit. Approximately 0.5 grams of intestinal content were diluted to about 500 vegetative cells of *C. guttulatus* per milliliter with sterilized distilled water. Then, 20 microliters of the suspension were added to 10 milliliter YPG (pH 1.5) medium supplemented with 100mg/L ampicillin. It was cultured on a 96-well culture plate with 100 microlitres of medium per well at 37°C with 10% CO_2_ for 2 hours. Wells with a single *C. guttulatus* vegetative cell were further cultured for 5 days before monoclonal cells of *C. guttulatus* were smeared and cultivated on solid YPG plate (pH 4.5) supplemented with 100 mg/L ampicilin at 37°C with 10% CO_2_. A single colony of *C. guttulatus* was transferred to liquid YPG medium (pH 4.5) and cultivated at 37°C on a orbital shaker (Zhichu, China) at a constant rotating speed of 200 rpm. Yeast multiplication was monitored using an automatic microbial growth curve analyser (Bioscreen, Finland) and an optical density scanner (Bug Lab, USA).Culture conditions were optimized by varying the medium pH and culture temperature.

### *C. guttulatus* identification

The morphology of *C. guttulatus* Zhejiang strain was observed under a light microscope with 400× magnification. Molecular identification was undertaken by PCR and gene sequencing. Two specific primer pairs for the small subunit (18S) and large subunit (26S) ribosomal RNA genes were synthesized according to Kurtzman CP (1998)^[23]^. The primer pairs were: 18S upper primer (TACGGTGAAACTGCGAATGG), 18S lower primer (GCTGATGACTTGCGCTTACT), 26S upper primer (GCATATCAATAAGCGGAGGAAAAG) and 26S lower primer (GGTCCGTGTTTCAAGACGG). The PCR conditions for the 18S DNA fragment were initial denaturation at 95 °C for 5 min, followed by 30 cycles of 94 °C denaturation for 45 sec, primer annealing at 55 °C for 45 sec, and extension at 72 °C for 90 sec. A final primer extension of 10 min at 72 °C completed the amplification process. The amplification for 26S was the same as for 18S except for extension at 72 °C for 120 sec during the PCR cycles. *C. guttulatus* vegetative cells were directly used as the template for PCR. The PCR products were examined and separated by 1% agarose gel electrophoresis. The target bands were purified by a gel extraction kit and sequenced by Sangon Biotech (Shanghai, China). The sequenced 18S and 26S gene fragments of the *C. guttulatus* Zhejiang isolate were submitted to Genbank with the Bankit procedure and blasted in the NCBI (National Center for Biotechnology Information, USA) database. Similar reference sequences were retrieved. The phylogenetic analysis was performed using the MegAlign program (DNAstar Inc., USA). The phylogenetic tree was constructed using the Clustal W method. Informations about *C. guttulatus* isolates and other yeast species used the construction of the phyogenetic trees are listed in Table 1.

### Animals

Specific-pathogen-free (SPF) rabbits were purchased from Pizhou Dongfang Rabbit Breeding Co., Ltd (Pizhou, China) and reared in our institute. Pre-weaning SPF rabbits were supplied with carrots and 0.5% milk powder in drinking water. No coccidia oocysts and *C. guttulatus* were detected in feces of these rabbits before use.

### Inoculation of *Cyniclomyces guttulatus* in rabbits and examination of yeast colonization in the gastrointestinal tract

Three groups of 20 day-old SPF rabbits (n=4) were orally inoculated with 1 × 10^6^, 1 × 10^7^ or 1 × 10^8^*C. guttulatus* vegetative cells per rabbit, and designated as G1, G2 and G3, respectively. Another group (G4) was not inoculated with the yeast and served as the control. Body weight, activities, appetite and excreta were recorded before and after yeast inoculation. Feces from all groups were collected daily and examined for yeast cells under a light microscope.

All rabbits were euthanized 18 days after inoculation. Contents and the mucous layer of the stomach, duodenum, jejunum, and ileum were collected, smeared and microscopically examined. Because the yeast was observed in the stomach (see Results), tissues from different regions of the stomach were immediately frozen for microscopic examination, and fixed in 10% formalin for preparation of paraffin sections for microscopic observation after staining with PAS (Periodic acid-schiff) and in 5% glutaraldehyde for preparation of electron-microscope sections for observation by transmission electron microscopy.

### Co-infection of *Cyniclomyces guttulatus* and *Eimeria intestinalis* in rabbits

Two groups (designated as CG and CG/EI) of 28 day-old SPF rabbits were orally inoculated with 4 ×10^7^ *C. guttulatus* cells per rabbit (n=4). After 14 days, CG/EI and another group (designated as EI, without *C. guttulatus* inoculation) were infected with 1 ×10^4^ sporulated oocysts of *E. intestinalis*. One group (designated as NON) served as un-infected control. Body weight, activities, appetite and excreta were recorded before and after inoculation. Feces were collected for coccidial oocysts and yeast cell counting. *E. intestinalis* oocysts in feces were counted between 10-16 days after infection using the McMaster method as described previously (Jeffers, 1975). Vegetative cells of *C. guttulatus* were counted 2 days before to 14 days after the infection of *E. intestinalis*. Briefly, one gram of feces was mashed with a glass stick and mixed with 60 milliliter of tap water. The fecal suspension was filtered through a 100-mesh sieve. *C. guttulatus* cells in the filtrate were counted in a haemocytometer under a light microscope with 100× magnification.

### Ethics statement

All experimental procedures were approved by the Zhejiang Academy of Agricultural Science Animal Ethics Committee (approval number, 20191904) and due attention was paid to the welfare of the animals. The rabbits were reared under stress-free environment, eliminating strong light and noise, with one rabbit per cage. Physical condition was monitored every day during all experimental procedures. Euthanasia was performed with an intra-cardiac pentobarbital overdose in accordance with the experiment design^[24]^.

### Prevalence survey

Fecal samples were collected from 253 healthy rabbits in four regions including Fuyang, Haining, Deqing and Wencheng (abbreviated names: FY, HN, ZX and WC) in Zhejiang province of China. The sample numbers from FY, HN, DQ and WC were respectively 50, 50 49 and 104. Among the surveyed rabbits, 66 were below 60 days old and 187 above 60 days old. The collected samples were stored at 4°C and examined within one week. The examination was performed as follows: 2 grams of feces were mixed with 60 milliliter of tap water. The mixture was filtered through a 100-meshsieve. Then, 100 microliter of filtrate was collected for wet mount examination under a light microscope with 100× magnification. Twenty fields were observed for each sample.

### Statistical analysis

Statistical analyses were performed for *C. guttulatus* cell counts, rabbit body weight, and *E. intestinalis* oocyst counts by GraphPad Prism 5.01. Data were expressed as mean±standard deviation, and the t-test was used to analyse differences between the mean values. Differences between groups with *p* values < 0.05 were considered statistically significant.

## Results

### A *Cyniclomyces guttulatus* Zhejiang strain was isolated and culture conditions optimized

A *C. guttulatus* Zhejiang strain was isolated from a diarrheic rabbit and cultivated in the YPG medium at low pH and identified by light microscopy. Microscopically, the vegetative cells of *C. guttulatus* were ellipsoid, colorless and about 20-50 μm in length and were occupied with two large vacuoles in the cytoplasm [Fig. 1 A&C]. *C. guttulatus* formed pseudohyphae when it was cultivated in a stationary culture station [Fig.1 B]. When it was cultivated with rotation, free vegetative cells were clearly visible. It could aerobically grow at a temperature range of 36 to 42 °C and a pH range of 1.5 to 4.5 in liquid YPG medium [Fig. 1 D&E]. The optimal culture medium pH was 4.5 [Fig.1 D]. The logarithmic growth phase was 24 to 60 hours at various temperatures and the pH value of 4.5 [Fig.1 E]. At the culture temperature of 40°C and pH 4.5, the cell density of *C. guttulatus* reached 4.62×10^7^±3.98×10^6^ cells per milliliter over 60 h from the initial culture density of 1×10^4^ cells per milliliter [Fig.1 E].

**Figure1.**
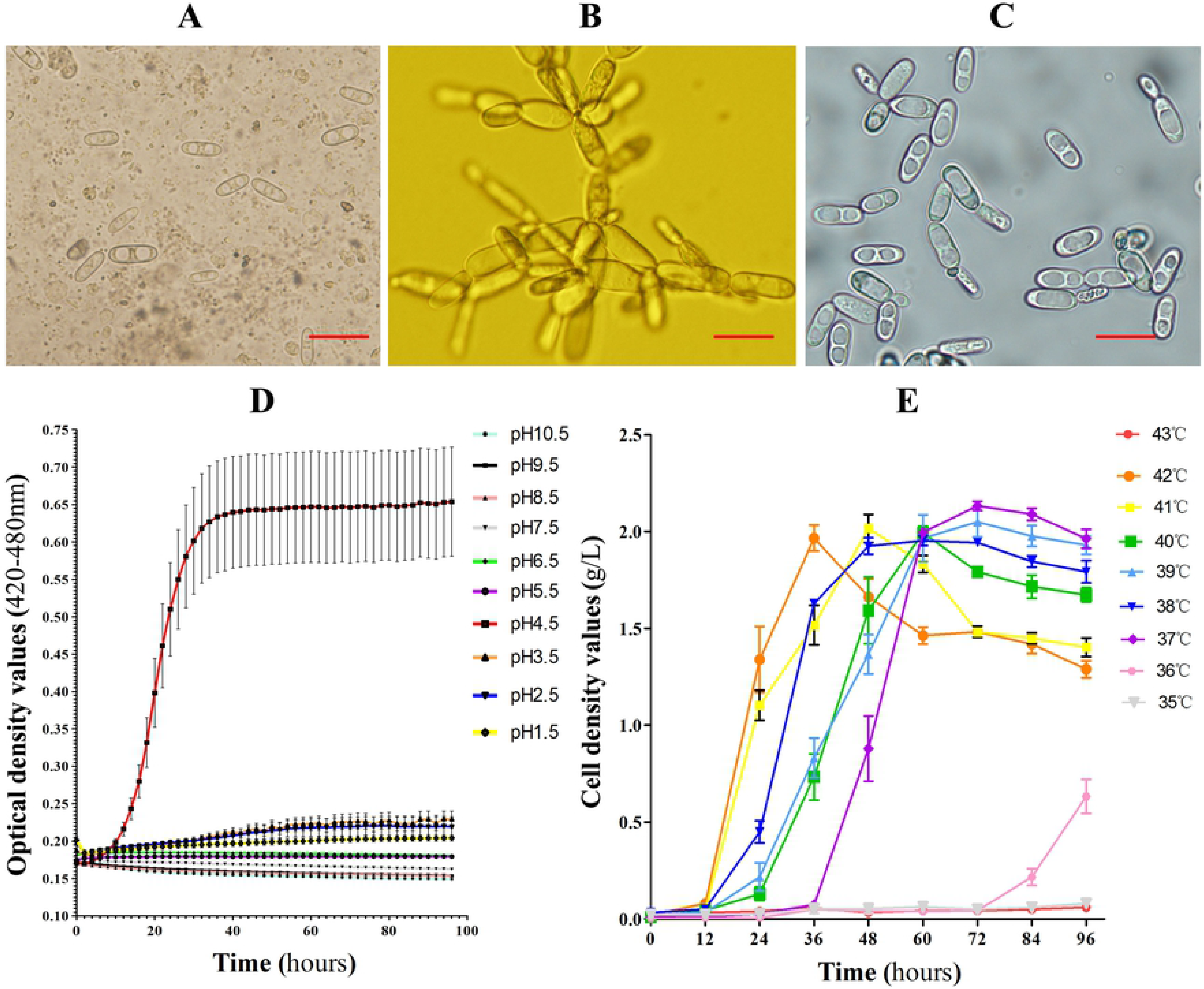

### The *C. guttulatus* Zhejiang strain showed a close relationship with reference strains originated from herbivores

The sequenced length for the 18S fragment of the *C. guttulatus* Zhejiang strain was 1559 bp. It showed 98% sequence identity and 100% coverage with that of the *C. guttulatus* NRRL Y-17561 strain reported in Genbank (accession number JQ698886.1)[Fig 2 A]. In the phylogenetic tree based on the 18S fragment, the Zhejiang strain clustered and formed a sister clade with the NRRL Y-17561 strain. The sequenced length for the 26S fragment of the *C. guttulatus* Zhejiang strain was 632 bp, and showed 100% (95% coverage), 100% (93% coverage), 97.2% (84% coverage), 96.6% (96% coverage) and 95.9% (95% coverage) identity with those of CGDPA-GP1, DPA-CGR1, Dog-1, DPA-CGD1 and NRRL Y-17561 strains, respectively. In the 26S phylogenetic tree, the Zhejiang strain was positioned in the same clade with CGDPA-GP1 and DPA-CGR1, and formed a sister clade with Dog-1, DPA-CGD1 and NRRL Y-17561 strains [Fig2 B]. According to data in the Genbank, *C. guttulatus* CGDPA-GP1 and DPA-CGR1 strains originated from herbivores, guinea pigs and rabbits, respectively, while *C. guttulatus* Dog-1 and DPA-CGD1 strains originated from the dog. Thus, the *C. guttulatus* Zhejiang strain showed a closer phylogenetic relationship with the strains originated from herbivores than those from the carnivore.

**Figure2.**
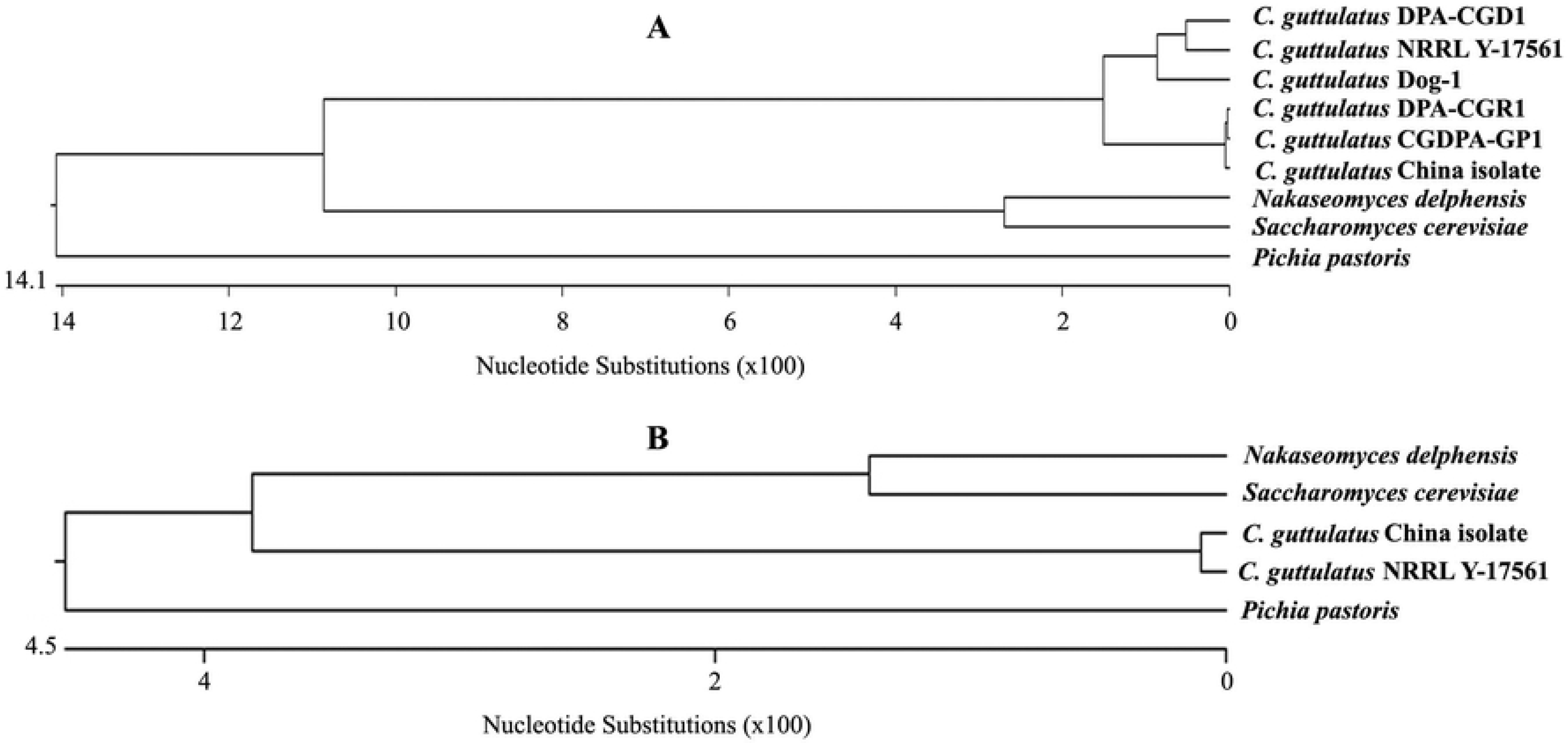

### *C. guttulatus* Zhejiang strain is non-pathogenic to healthy rabbits

To study the pathogenicity of *C. guttulatus*, we inoculated SPF rabbits with a large dose of yeast cells. Two days after inoculation, vegetative cells of *C. guttulatus* were detectable in rabbit feces. None of the rabbits inoculated with 1×10^6^ to 10^8^ of *C. guttulatus* showed clinical signs of illness. Interestingly, *C. guttulatus*-innoculated groups (G1-G3) had less feed waste than the control group (G4) [Fig3 B&C]. Mean body weight of inoculated groups was slightly higher than that of the control group although the difference was not statistically significant (*p*>0.05) [Fig3. A]. Autopsy showed no macroscopic or microscopic lesions in the gastrointestinal tract of the inoculated rabbits despite a large number of *C. guttulatus* cells in the gastric and intestinal contents. Especially, a thick layer of *C. guttulatus* cells colonized the gastric mucosa [Fig3 D-F]. PAS-stained gastric tissue sections showed a dense layer of saccharides on the gastric mucosa and also on the cell wall of *C. guttulatus* [Fig3 G]. The *C. guttulatus* cells probably attached to the stomach mucosa through these filamentous saccharides as shown by transmission electron microscopy [Fig3 H&I]. Our findings indicate that the *C. guttulatus* Zhejiang strain colonizes on the gastric mucosa, but it is non-pathogenic to healthy rabbits.

**Figure3.**
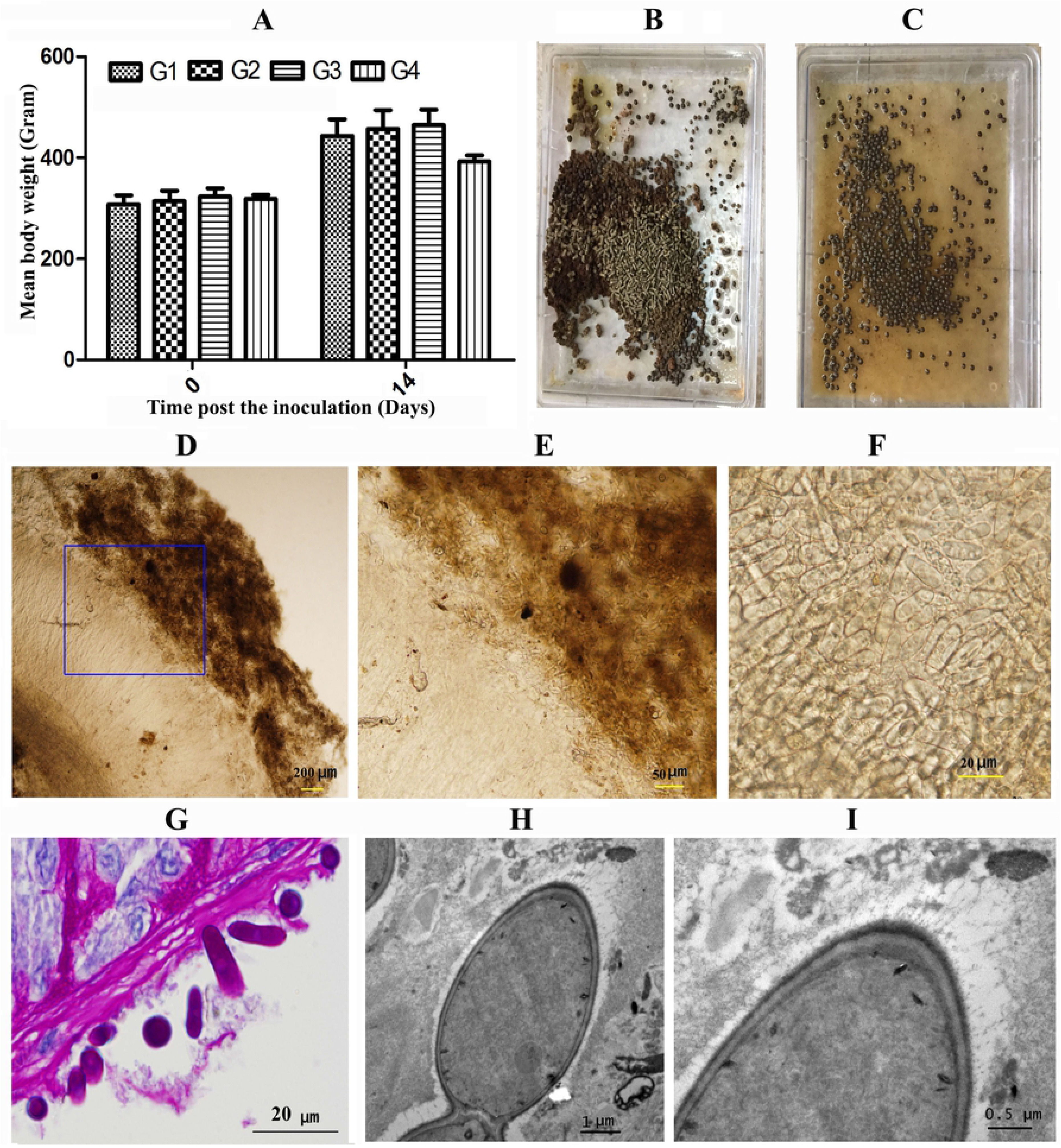

### *C. guttulatus* is an opportunistic pathogen in rabbits infected with *Eimeria intestinalis*

We investigated whether *C. guttulatus* could be an opportunistic pathogen in rabbits infected with coccidia. Rabbits pre-inoculated with *C. guttulatus* (CG/EI group) developed more severe illness and intestinal lesions following the infection of *E. intestinalis* than those non-inoculated with *C. guttulatus* (EI group). Compared with the EI group, more severe diarrhea, loss of appetite and constipation were observed in the CG/EI group. The number of *C. guttulatus* vegetative cells in feces of the CG/EI group was significantly higher than that of the CG group (*p*<0.05) on 9 and 10 days post *E. intestinalis* infection [Fig4 A]; the mean cells per gram of feces in the CG/EI group were 1.22 × 107 ± 1.38 × 106 on day 9 and 8.55 × 106 ± 5.52 × 10^5^ on day 10, 4.7 fold and 3.2 fold higher than those of the EI group, respectively. In contrast, coccidian reproduction was lower in the co-inoculation rabbits than the rabbits inoculated with *E. intestinalis* alone, as shown by a markedly lower oocyst output in the CG/EI group than in the EI group; the total fecal oocyst count per rabbit on day 10 was 2.87×10^9^±8.13×10^7^ in the co-inoculation group, compared with 4.57×10^9^±6.83×10^7^ in the group without yeast inoculation [Fig4 B]. In addition, the peak oocyst excretion of the CG/EI group was also lower than that of the EI group (1.04×10^9^±2.45×10^7^ on day 11 and 1.61×10^9^±3.43×10^7^ on day 13). Fecal oocyst excretion in the CG/EI group both peaked and cleared earlier than that in the EI group [Fig4 B]. Thus, *C. guttulatus* proliferation was enhanced by *E. intestinalis* infection, while *E. intestinalis* reproduction was suppressed by *C. guttulatus*. In addition, one rabbit of the CG/EI group died on the 10th day post infection. The dead rabbit had disseminated hemorrhage and nodules in the lower jejunum and ileum and a large number of *E. intestinalis* oocysts and *C. guttulatus* vegetative cells were detected in the small intestine [Fig4 D&E]. In addition, mean body weight of the CG/EI group was slightly lower than the CG group 11 to 17 days post infection [Fig4 C]. In summary, rabbits inoculated with both *E. intestinalis* and *C. guttulatus* developed more severe clinical signs and intestinal lesions (and one death) than the rabbits inoculated with *E. intestinalis* only, associated with greater *C. guttulatus* and lower *E. intestinalis* output. The findings suggest *C. guttulatus* may be an opportunistic pathogen to rabbits with coccidia.

**Figure4.**
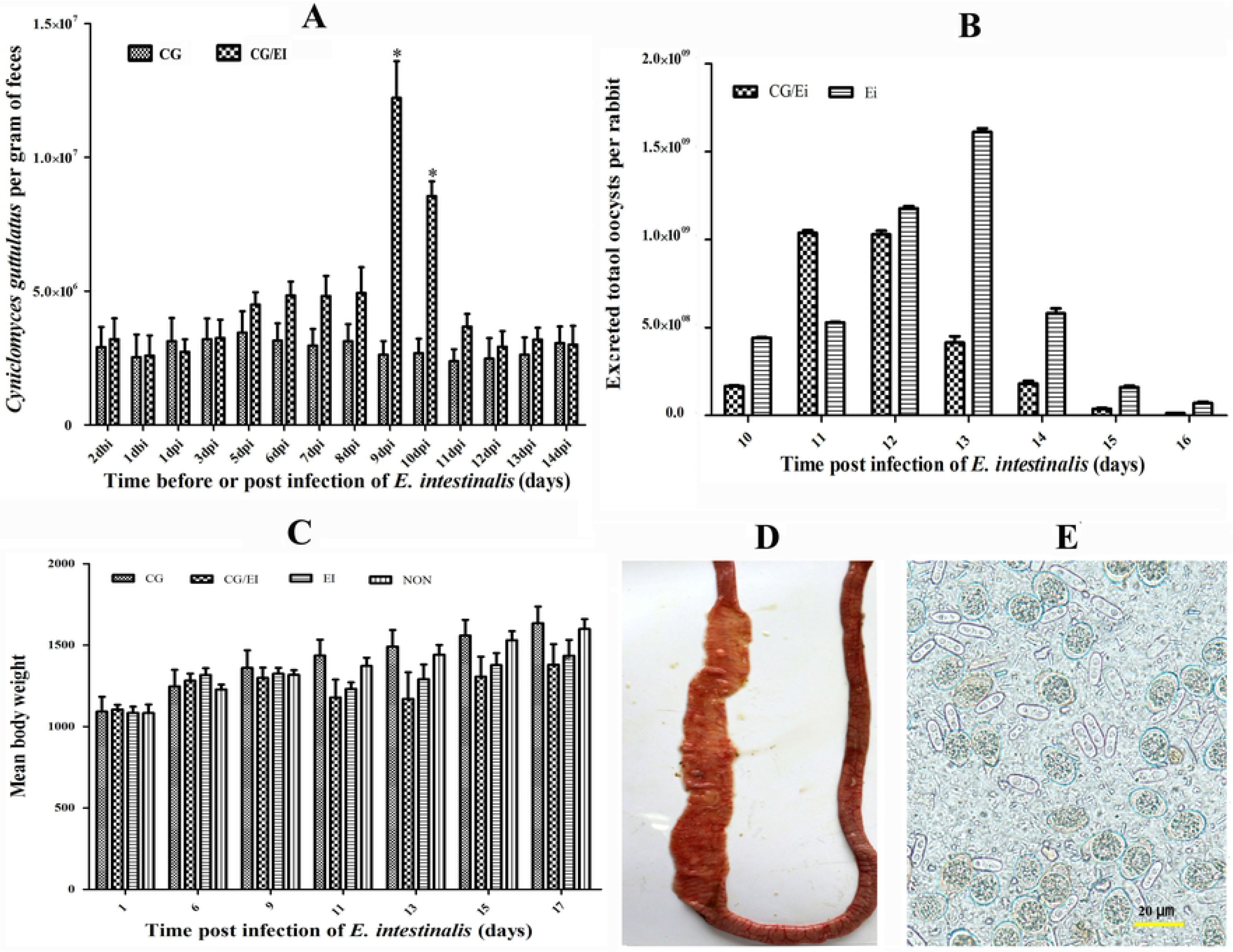

### *C. guttulatus* was highly prevalent in rabbits

We conducted a survey of rabbits carrying *C. guttulatus* by analyzing *C. guttulatus* cells in feces. *C. guttulatus* was detected in 210 of 253 fecal samples from rabbits above 30 days old in Zhejiang province, a positive rate of 83%. Of the surveyed rabbits, the positive rate for rabbits above 60 days old was 69.7% (46/66) and for rabbits less than 60 days old 87.7% (164/187). The number of rabbits with a high *C. guttulatus* load (>100 cells per microscopic field at 200x magnification) were 59 (23.3%), consisting of 4 rabbits less than 60 days old and 55 rabbits above 60 days old (6.1% and 29.4% of the age group, respectively). Thus, *C. guttulatus* was highly prevalent in healthy rabbits, especially in older rabbits.

## Discussion

*Cyniclomyces guttulatus* is a monotypic yeast genus of the Saccharomycetaceae family, and inhabits the gastrointestinal tract of many animal species including rabbits, dogs and guinea pigs^[17,22,25]^. *C. guttulatus* was first described more than 60 years ago, and is commonly found in rabbit feces. A large number of *C. guttulatus* cells are often found in feces of diarrheic rabbits, but it is unknown whether *C. guttulatus* causes diarrhea in rabbits. We isolated a *C. guttulatus* Zhejiang strain from a rabbit with severe diarrhea. At optimized culture pH and temperature, a single clone was expanded and studied for its pathogenicity in rabbits. We demonstrated that the *C. guttulatus* Zhejiang strain is not a primary pathogen causing diarrhea in healthy SPF rabbits. Inoculation of as high as 1 × 10^8^ vegetative cells per rabbit did not result in any clinical signs of illness or gastrointestinal lesions. This is consistent with previously reported studies, where rabbits orally or intravenously inoculated with *C. guttulatus* isolates showed no clinical signs of illness^[16,17,26]^. Unexpectedly, we found that rabbits inoculated with *C. guttulatus* showed better performance in terms of body weight gain and food intake. *C. guttulatus* seems a probiotic microorganism in rabbits, especially in weanling rabbits. However, the findings of massive vegetative cells of *C. guttulatus* commonly seen in feces of rabbits with diarrhea suggest it could be a causative microorganism of GI tract disturbance.

Some authors proposed *C. guttulatus* could be an opportunistic pathogen or play a co-causative role in diarrhea of its host, based on indirect evidence that the antifungal agent, nystatin was effective in the treatment of some diarrheic cases^[19–22]^. In our study, *C. guttulatus* was proved an opportunist through co-infection with the coccidian species *E. intestinalis*, a parasite causing diarrhea and intestinal lesions in rabbits. Mortality and more severe clinical signs and intestinal lesions were observed in rabbits co-infected with *C. guttulatus* and *E. intestinalis* than in rabbits inoculated with *E. intestinalis* only. Compared with rabbits inoculated with *C. guttulatus* alone, vegetative cells of *C. guttulatus* more prolificly multiplied in the co-infection group, peaking at 9-10 days post *E. intestinalis* infection. This time period is consistent with that of intestinal lesions of *E. intestinalis* infection^[27,28]^. The massive multiplication of *C. guttulatus* was probably a consequence of altered gastrointestinal environment in the rabbits from *E. intestinalis* infection. Compared with rabbits with *E. intestinalis* infection alone, *E. intestinalis* oocyst excretion in the co-infected rabbits decreased by 37%. We speculate that the rapid and massive multiplication of *C. guttulatus* vegetative cells could contribute to the severe symptoms in the co-infected rabbits, and the massive multiplication of *C. guttulatus* also limits the reproduction of *E. intestinalis*.

In addition, our epidemiological survey showed *C. guttulatus* is prevalent in rabbits in Zhejiang, China. The positive rate in rabbits was as high as 83%. This was significantly higher than that in other host animals such as dogs in which a prevalence of 14-21% was reported^[20,21,29]^. Coccidia are also highly prevalent in rabbits. According to a previous study, the overall prevalence of rabbit coccidia is 41.9% in China, and as high as 70% in some regions^[12]^. So, *C. guttulatus* may contribute to the morbidity and mortality of rabbits with coccidiosis.

In summary, *C. guttulatus* as a commensal yeast is very common in rabbits. It is usually not pathogenic in healthy rabbits and seems a probiotic microorganism in rabbits, but it could become an opportunistic pathogen when the gastrointestinal environment is altered by enteric pathogens such as coccidia. Considering the high prevalence of both *C. guttulatus* and coccidia in rabbits, the potential harm of *C. guttulatus* to the rabbit industry warrants attention and further study.

## Acknowledgements

This research work was supported by the National Natural Science Foundation of China (Project number 31101810), Project of Zhejiang Province Public Welfare Technology Application Research Project (Project number 2017C32037), China Agricultural Research System (Project number nycytx-44-3-2) and Key Research and Development Plan of Zhejiang Province (Project numbers 2016C02054-10, 2019C02052).

